# Evolinc: a comparative transcriptomics and genomics pipeline for quickly identifying sequence conserved lincRNAs for functional analysis

**DOI:** 10.1101/110148

**Authors:** Andrew D. L. Nelson, Upendra K. Devisetty, Kyle Palos, Asher K. Haug-Baltzell, Eric Lyons, Mark A. Beilstein

## Abstract

Long intergenic non-coding RNAs (lincRNAs) are an abundant and functionally diverse class of eukaryotic transcripts. Reported lincRNA repertoires in mammals vary, but are commonly in the thousands to tens of thousands of transcripts, covering ~90% of the genome. In addition to elucidating function, there is particular interest in understanding the origin and evolution of lincRNAs. Aside from mammals, lincRNA populations have been sparsely sampled, precluding evolutionary analyses focused on lincRNA emergence and persistence. Here we present Evolinc, a two-module pipeline designed to facilitate lincRNA discovery and characterize aspects of lincRNA evolution. The first module (Evolinc-I) is a lincRNA identification workflow that also facilitates downstream differential expression analysis and genome browser visualization of identified lincRNAs. The second module (Evolinc-II) is a genomic and transcriptomic comparative analyses workflow that determines the phylogenetic depth to which a lincRNA locus is conserved within a user-defined group of related species. Evolinc-II builds families of homologous lincRNA loci, aligns constituent sequences, infers gene trees, and then uses gene tree / species tree reconciliation to reconstruct evolutionary processes such as gain, loss, or duplication of the locus. Here we demonstrate that Evolinc-I is agnostic to target organism by validating against previously annotated Arabidopsis and human lincRNA data. Using Evolinc-II, we examine ways in which conservation can rapidly be used to winnow down large lincRNA datasets to a small set of candidates for functional analysis. Finally, we show how Evolinc-II can be used to recover the evolutionary history of a known lincRNA, the human telomerase RNA (TERC). The analyses revealed unexpected duplication events as well as the loss and subsequent acquisition of a novel TERC locus in the lineage leading to mice and rats. The Evolinc pipeline is currently integrated in CyVerse’s Discovery Environment and is free to use by researchers.

## Introduction

A large, and in some cases predominant, proportion of eukaryotic transcriptomes are composed of long non-coding RNAs (lncRNAs) (Guttman et al., 2009; Cabili et al., 2011; Wang et al., 2015; Liu et al., 2012). LncRNAs are longer than 200 nucleotides (nt) and exhibit low protein-coding potential (non-coding). While some transcripts identified from RNA-seq are likely the result of aberrant transcription or miss-assembly, others are bona fide lincRNAs with various roles (see [Wang and Chang, 2011; Ulitsky and Bartel, 2013] for a review of lncRNA functions). To help factor out transcriptional “noise”, additional characteristics are used to delineate lncRNA. These additional characteristics focus on factors such as reproducible identification between experiments, degree of expression, and number of exons (Derrien et al., 2012). In general, lncRNAs display poor sequence conservation between closely related species, are expressed at lower levels than protein-coding genes, and largely lack functional data.

The function of any particular lncRNA is likely to influence its evolution. One means of inferring that a transcript is a functional lncRNA and not an artefact is the degree of conservation we observe at that locus between two or more species. This conservation can be observed at the sequence, positional, and transcriptional level (Ulitsky, 2016). Comparative approaches using these criteria of conservation have been successful to varying degrees in different eukaryotic lineages. To simplify matters, these approaches typically focus on long intergenic non-coding RNAs (lincRNAs), as their evolution is not constrained by overlap with protein-coding genes. In vertebrates, lincRNA homologs have been identified in species that diverged some 400 million years ago (MYA), whereas in plants lincRNAs homologs are primarily restricted to species that diverged < 100 MYA (Ulitsky et al., 2011; Necsulea et al., 2014; Nelson et al., 2016; Liu et al., 2012; Li et al., 2014; Zhang et al., 2014; Mohammadin et al., 2015). Importantly, the conserved function of a handful of these lincRNAs have been experimentally verified *in vivo* (Hawkes et al., 2016; Migeon et al., 1999; Quinn et al., 2016).

One major factor inhibiting informative comparative genomics analyses of lincRNAs is the lack of robust sampling and user-friendly analytical tools. Here we present Evolinc, a lincRNA identification and comparative analysis pipeline. The goal of Evolinc is to rapidly and reproducibly identify candidate lincRNA loci, and examine their genomic and transcriptomic conservation. Evolinc relies on high quality RNA-seq data to annotate putative lincRNA loci across the target genome. It is designed to utilize cyberinfrastructure such as the CyVerse Discovery Environment (DE), thereby alleviating the computing demands associated with transcriptome assembly (Merchant et al., 2016). The pipeline is divided into two modules. The first module, Evolinc-I, identifies putative lincRNA loci, and provides output files that can be used for analyses of differential expression, as well as visualization of genomic location using the EPIC-CoGe genome browser (Lyons et al., 2014). The second module, Evolinc-II, is a suite of tools that allows users to identify regions of conservation within a candidate lincRNA, assess the extent to which a lincRNA is conserved in the genomes and transcriptomes of related species, and explore patterns of lincRNA evolution. We demonstrate the versatility of Evolinc on both large and small datasets, and explore the evolution of lincRNAs from both plant and animal lineages.

## Materials and Methods

In this section we describe how the two modules of Evolinc (I and II) work, as well as describing the data generated by each.

### Evolinc-I: LincRNA identification

Evolinc-I minimally requires the following input data: a set of assembled and merged transcripts from Cuffmerge or Cuffcompare (Trapnell et al., 2010) in gene transfer format (GTF), a reference genome (FASTA), and a reference genome annotation (GFF). Evolinc places all unknown or novel gene variants, including the various overlapping class codes assigned by Cufflinks (Trapnell et al., 2010) into a new file for downstream filtering. From these transcripts, only those longer than 200 nt are kept for further analysis. Transcripts with high protein-coding potential are removed using two metrics: 1) open reading frames (ORF) encoding a protein > 100 amino acids, and 2) similarity to the uniprot protein database (based on a 1E-5 threshold). Filtering by these two metrics is carried out by Transdecoder (https://transdecoder.github.io/) with the BLASTp step included. These analyses yield a set of transcripts that fulfill the basic requirements of lncRNAs.

The role of transposable elements (TEs) in the emergence and function of lncRNAs is an active topic of inquiry (Wang et al., 2017; Kapusta et al., 2013). To facilitate these studies, Evolinc allows the user to separate lncRNAs bearing similarity to TEs into a separate FASTA file. This is performed by BLASTn (Altschul et al., 1990), with the above lncRNAs as query against a user provided TE database (in FASTA format). We considered lncRNAs that exceeded a bit score value of 200 and an E-value threshold of 1E-20 to be TE-derived. These stringent thresholds remove TE-derived lncRNAs with high similarity to TEs, but allow for retention of lncRNAs with only weak similarity to TEs, perhaps reflecting older TE integration events or TE exaptation events (Johnson and Guigó, 2014). To thoroughly identify TE-derived lncRNAs, we suggest building the TE database from as many closely related and relevant species as possible. Along with these sequence files, BED files are also generated for these TE-derived lncRNAs for further analysis by the user, but are not used in downstream analyses by Evolinc-I.

Candidate lncRNAs are next compared against reference annotation files to determine any overlap with known genes. Some reference annotations distinguish between protein-coding and other genes (lncRNAs, pseudogenes, etc). If this style of reference annotation is available, we suggest running Evolinc-I twice, once with an annotation file containing only protein-coding genes and once with all known genes. This is a simple way to distinguish between the identification of novel putative lncRNAs and known (annotated) lncRNAs. We also recommend using an annotation file that contains 5’ and 3’ UTRs where possible. If this is unknown, the genome coordinates within the reference annotation file should be manually adjusted to include additional sequence on either end of known genes (i.e., 500bp). This number can be adjusted to adhere to community-specific length parameters for intergenic space. Evolinc-I will identify lncRNAs that overlap with the coordinates of a known gene and then sort into groups of overlapping lncRNAs based on directionality: sense and antisense-overlapping lncRNA transcripts (SOT and AOT, respectively). Keep in mind that in order for these inferences to be made, either strand-specific RNA-sequencing must be performed or the lncRNA must be multi-exonic. Sequence FASTA and BED files for each group of overlapping lncRNAs are generated by Evolinc-I for the user to inspect. Demographic data are also generated for each of these lncRNA types (explained further below).

LncRNAs that do not overlap with known genes and have passed all other filters are considered (putative) lincRNAs. Evolinc-I also deals with optional input data that may increase the confidence in the validity of particular candidate lincRNAs. For example, when users provide transcription start site coordinates (in BED format), Evolinc-I identifies lincRNAs in which the 5’ end of the first exon is within 100bp of any transcriptional start site (TSS). LincRNAs with TSS are annotated as “CAGE_PLUS” in the FASTA sequence file (lincRNAs.FASTA), and the identity of such lincRNAs is recorded in the final summary table (Final_summary_table.tsv). Optionally, Evolinc-I identified lincRNAs (termed Evolinc-lincRNAs) can also be tested against a set of user-defined lincRNAs that are not found in the reference annotation (i.e., an in-house set of lincRNAs not included in the genome annotation files or if annotation for a particular system lags behind lincRNA identification). When the coordinates for a set of such lincRNA loci are provided in general feature format (GFF), Evolinc-I will use these data to determine if any putative Evolinc-lincRNAs are overlapping. These loci are appended with “_overlapping_known_lncRNA” in the lincRNA.FASTA file. The identity of the overlapping (known lncRNA) is listed for each Evolinc-lincRNA in the final summary table.

### Output from Evolinc-I

Evolinc-I generates a sequence file and BED file for TE-derived lncRNAs, AOT or SOT lncRNAs, and intergenic lncRNAs (lincRNAs). An updated genome annotation file is created, appending only the lincRNA loci to the user-supplied reference annotation file. This file can then be used with differential expression analysis programs such as DESeq2 or edgeR (Anders and Huber, 2010; Robinson and Oshlack, 2010). In addition, two types of demographic outputs are generated. For SOT, AOT, and lincRNAs, a report is created that describes the total number of transcripts identified for each class (isoforms and unique loci), GC content, minimum, maximum, and average length. For lincRNAs only, a final summary table is generated with the length and number of exons for each lincRNA, as well as TSS support and the ID of any overlapping, previously curated lincRNA. The Evolinc-I workflow is shown in Figure 1A.

**Figure 1.**
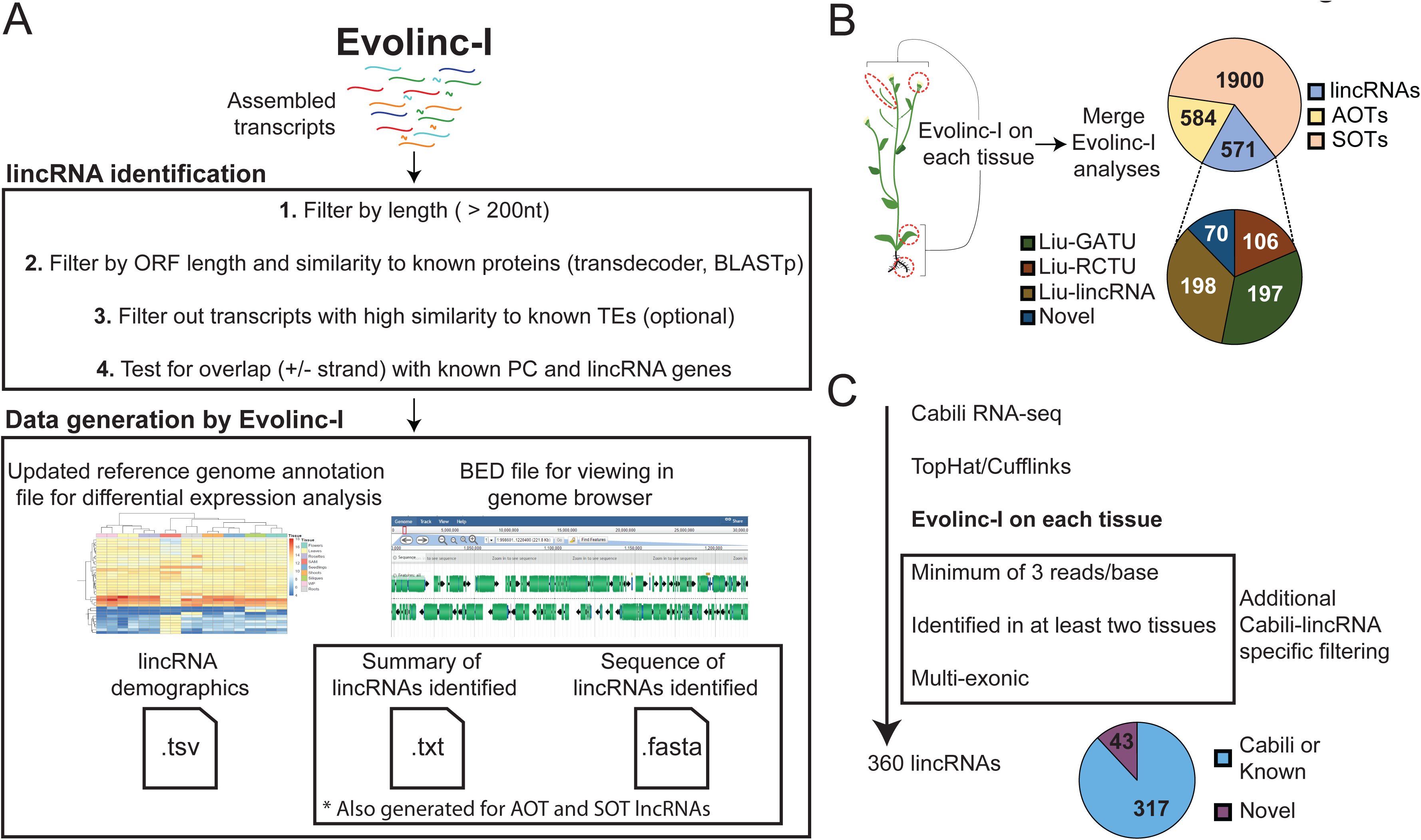
Schematic representation of the Evolinc-I workflow and validation. **(A)** Evolinc-I takes assembled transcripts as input and then filters over several steps (1-4). Evolinc generates several output files detailed in the materials and methods. **(B)** Evolincvalidation on RNA-seq data from Liu et al. (2012). Four tissues were sequenced by Liu et al, as indicated by the red circles, including (from top to bottom) flowers, siliques, leaves, and roots. Assembled transcripts were fed through Evolinc-I to identify lincRNAs, Antisense Overlapping Transcripts (AOTs), and Sense Overlapping Transcripts (SOTs). A reconciliation was performed between the Evolinc-I identified lincRNAs and the Liu et al. dataset. Gene associated transcriptional unit (GATU) amd repeat containing transcriptional unit (RCTU) terminology comes from Liu et al. (2012). **(C)** Evolinc validation of Cabili et al. (2011) RNA-seq data. RNA-seq data was assembled and then filtered through additional Cabili-specific parameters (shown in box). Shown in the pie chart are the Evolinc-identified lincRNAs that correspond to Cabili et al. or are novel.

### Identifying lincRNA conservation with Evolinc-II

Evolinc-II minimally requires the following input data: a FASTA file of lincRNA sequences, FASTA file(s) of all genomes to be interrogated, and a single column text file with all species listed in order of phylogenetic relatedness to the query species (example and further elaboration on the species list in File S1). Many of these genomes can be acquired from CoGe (www.genomevolution.org), and lincRNA sequences can be obtained from either the output of Evolinc-I or from another source. The number, relationship, and divergence times of the genomes chosen will depend on the hypotheses the user intends to test. To determine the transcriptional status of lincRNA homologs across a group of species, Evolinc-II can optionally incorporate genome annotation files (GTF) and known lincRNA datasets from target species in FASTA format. In addition, Evolinc-II can incorporate motif and structure data, in BED format, to highlight any potential overlap between conserved regions and user-supplied locus information.

Evolinc-II starts by performing a series of reciprocal BLASTn searches against provided target genomes, using a user-defined set of lincRNAs as query and user chosen E-value cutoff. We suggest starting at an E-value cutoff of 1E-20 because we found that, across 10 Brassicaceae genomes, and independently between human, orangutan and mouse, this value was optimal for recovering reciprocal and syntenic sequence homologs (Nelson et al., 2016). While 1E-20 represents a starting point for these analyses, lincRNA homolog recovery relies on a variety of factors (i.e., background mutation rate, genome stability, evolutionary distance of species / taxa being analyzed) that could affect the E-value cutoff most likely to return homologous loci among related genomes. Thus, we recommend “calibrating” Evolinc-II using at least three genomes of varying divergence times from the query species before including a larger (> 3) set of genomes. After BLASTn, the top blast hit (TBH) in each species for the query lincRNA is identified. Multiple, non-redundant hits falling within the same genomic region, which is likely to occur when the query lincRNA is multi-exonic, are merged as a single TBH. Sequence for all TBHs are then used as query in reciprocal BLAST searches (see below). For researchers interested in testing the evolutionary origins of specific lincRNAs, all BLAST hits that passed the E-value cutoff are retained. However, to reduce computing time, subsequent analyses are confined to TBHs. For query lincRNAs for which a TBH is not identified in the first iteration (i.e., did not pass the E-value cutoff), the query lincRNA is divided non-overlapping segments and each segment is used as query in a second set of BLAST searches.

TBHs from each species included in the analysis are then used as query sequences in a reciprocal BLAST against the genome of the species whose lincRNA library was used in the original query. For a locus to be considered homologous to the original query lincRNA locus, both loci must be identified as the TBH. This is especially useful when performing searches using a low E-value cutoff value, as false positives are reduced dramatically. TBHs that pass the reciprocity test are appended with “Homolog” in the final FASTA sequence alignment file (“query_lincRNA_1”_alignment.FASTA).

As TBHs from each target genome are identified, they are scanned for overlap against optional genome reference annotation datasets (GFF) and known lincRNA files (FASTA). The identifier number (ID) of all TBHs with overlap against these two datasets is appended with either “Known_gene” or “Known_lincRNA”. The identity of the overlapping gene is retained in the final summary table (final_summary_table.tsv) as well as in each FASTA sequence alignment file (see below). Many genes and almost all lincRNAs are annotated based on transcriptional evidence. Thus, this is a simple way of determining if a query lincRNA corresponds to a locus with evidence of transcription in another species. In addition, when working with a poorly annotated genome, comparing against well-annotated species can provide additional levels of information about the putative function of query lincRNAs. For example, if the homologous locus of a query lincRNA overlaps a protein-coding gene in that species, it could indicate that the query lincRNA is a protein-coding gene, or a pseudogene.

All TBH sequences for a given query lincRNA are then clustered into a family. For example, an Evolinc-II analysis that queries ten lincRNAs across a set of target genomes will result in ten lincRNA families, populated with the TBH from each target genome. Genomes that do not return a TBH at the specified E-value cutoff (from either full-length or segmented searches), or whose TBH does not pass the reciprocity test, will not be represented in the family. These lincRNA families are then batch aligned using MAFFT under default settings (Katoh and Standley, 2013). The alignment file for each lincRNA family can be downloaded into a sequence viewer. Evolinc-II will also infer phylogeny from the sequence alignment using RAxML (Stamatakis, 2014). Gene trees are reconciled with a user-provided species tree, in Newick format, using Notung (Durand et al., 2006). This latter analysis pinpoints duplication and loss events that may have occurred during the evolution of the lincRNA locus. The Notung reconciled tree is available to view in PNG format within the CyVerse DE. Duplication and loss events are denoted by a red D or L, respectively (Example in Figure S4). The Evolinc-II workflow is shown in Figure 2A.

**Figure 2.**
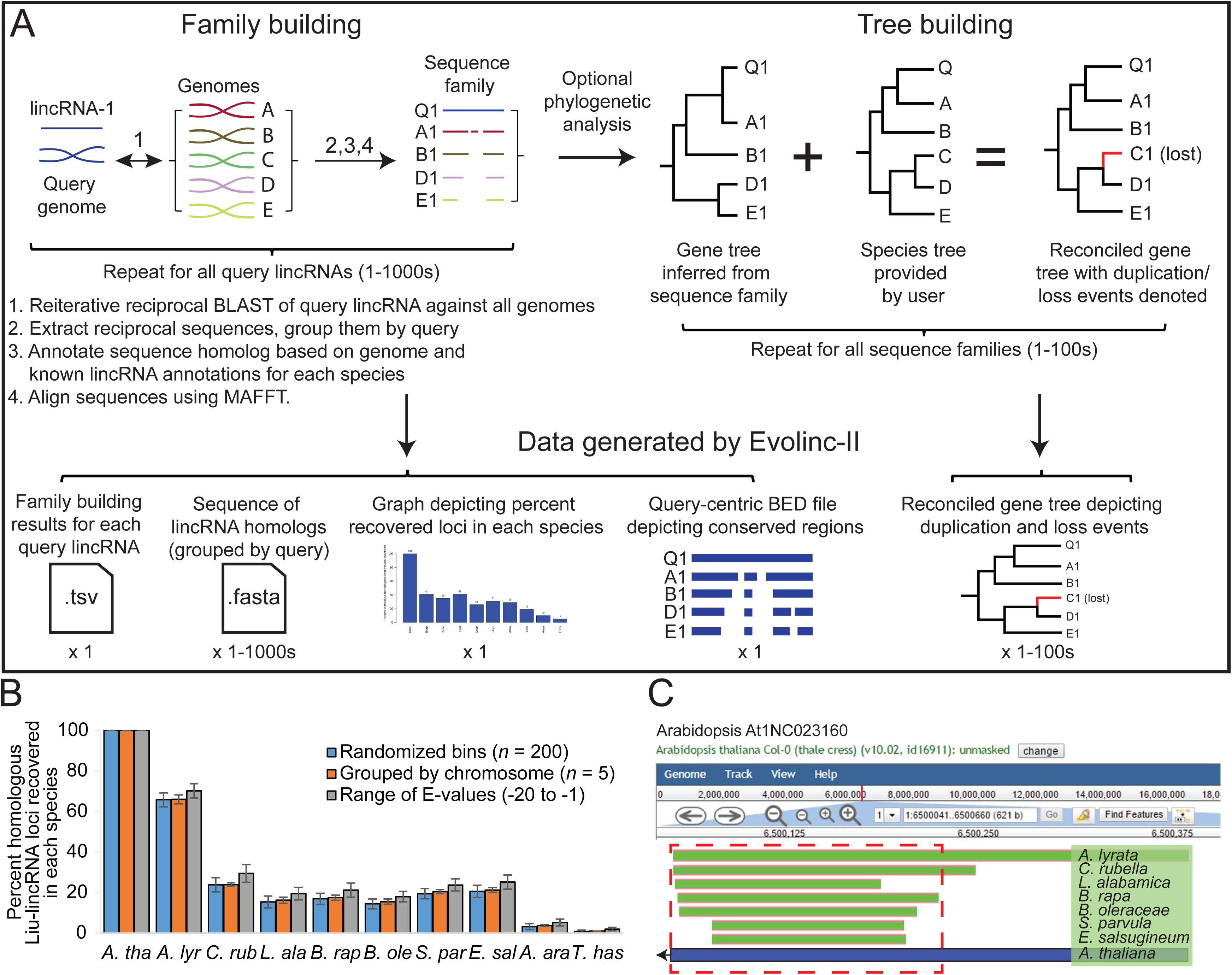
Schematic representation of the Evolinc-II workflow and validation of Liu-lincRNA and Evolinc-identified lincRNAs. **(A)** Evolinc-II uses lincRNAs as a query in reciprocal BLAST analyses against any number of genomes. Sequences that match the filters (see Materials and Methods) are grouped into families of sequences based on the query lincRNA. Each sequence homolog is annotated based on information about the locus in question, such as expression or overlap with known gene or lincRNA. Sequences are aligned to highlight conserved regions and prepare the sequence families for the optional phylogenetic steps. These can realistically be performed on thousands to tens of thousands of query lincRNAs. If selected, gene trees are inferred for each sequence family with RAxML and then reconciled to the known species tree using Notung 2.0. Notung delineates gene loss and duplication events by marking the output tree (png) with a D (duplication) and blue line, or L (loss) and a red line. As the RAxML step is computationally intensive, we suggest limiting the number of sequence families this is performed in an analysis or expect long compute times. Data files generated by Evolinc-II are described in the Materials and Methods. **(B)** Validation of Evolinc-II by repeating the Liu-lincRNA dataset in three different ways. The ~5400 Liu-lincRNAs were randomly divided into 200 sequence bins (blue bar), each bin was run through Evolinc-II (total number of runs = 27), and then the results were averaged, with standard deviation denoted. In the second analysis, the Liu-lincRNAs were divided based on chromosome, and then each set of Liu-lincRNAs (five groups) were run through Evolinc-II separately. Lastly, all Liu-lincRNAs were run through Evolinc using different BLAST E-values (E-1, -5, -10, -15, -20), then the results were averaged. Bars represent the percent of Liu-lincRNAs for which sequence homologs were identified across Brassicaceae. **(C)** Genome browser visualization of the At1NC023160 locus and its conservation in other Brassicaceae. Regions of the Arabidopsis locus that Evolinc-II identified to be conserved are shown in green, with species of origin listed to the right. The blue bar indicates the length of the locus in Arabidopsis, with the arrow indicating direction of transcription. The region of the locus selected for structural prediction is shown in the red dashed box.

### Output from Evolinc-II

Evolinc-II generates sequence files containing lincRNA families with all identified sequence homologs from the user-defined target genomes. In addition, a summary statistics table is generated of identified lincRNA loci based on depth of conservation and overlapping features (e.g., genes, lincRNAs, or other user defined annotations). In addition, the identity of overlapping features (e.g., gene, known lincRNAs) in each genome for which a sequence homolog was identified are listed (Shown for the Liu-lincRNAs in File S3). To visualize conserved regions of all query lincRNAs, a query-centric BED file is generated that is ready for import into any genome browser. An example using the genome browser embedded within CoGe (Tang and Lyons, 2012) is shown below (Figure 2C). Following phylogenetic analysis, a reconciled gene tree is produced with predicted duplication and loss events indicated. Lastly, to provide the user with a broad picture of lincRNA conservation within their sample set, a bar graph is produced that indicates the number and percent of recovered sequence homologs in each species (Figure S2A).

### Data and software availability

All genomes used in this work, including version and source, are listed in File S1. The accession number of all SRAs used in this work, including project ID, TopHat (Kim et al., 2013) read mapping rate, and total reads mapped for each SRA are shown in File S1. Genomic coordinates for lincRNAs identified by Evolinc-I are listed by species in BED/GFF format in File S2. Novel lincRNAs have also been deposited within the CoGe environment as tracks for genome browsing (Links found in File S2). Evolinc is available as two apps (Evolinc-I and Evolinc-II) in CyVerse’s DE (https://de.cyverse.org/de/), for which a tutorial and sample data are available (https://wiki.cyverse.org/wiki/display/TUT/Evolinc+in+the+Discovery+Environment). Evolinc is also available as self-contained Docker images (https://hub.docker.com/r/cyverse/evolinc-i/; and https://hub.docker.com/r/cyverse/evolinc-ii/) for use in a Linux or Mac OSX command-line environment. The code for Evolinc is available as a github repository (https://github.com/Evolinc). Both Evolinc tools make use of several open source tools, such as BLAST for sequence comparisons (Altschul et al., 1990), Cufflinks (Trapnell et al., 2010) for GFF to FASTA conversion, Bedtools (Quinlan and Hall, 2010) for sequence intersect comparisons, MAFFT (Katoh and Standley, 2013) for sequence alignment, RAxML (Stamatakis, 2014) for inferring phylogeny, Notung (Durand et al., 2006) for reconciling gene and species trees, and python, perl, and R for file manipulation and data reporting.

### RNA-seq read mapping and transcript assembly

SRA files were uploaded directly into CyVerse DE from (http://www.ncbi.nlm.nih.gov/sra) by using the “Import from URL” option. All further read processing was performed using applications within DE. Briefly, uncompressed paired end reads were trimmed (5 nt from 5’ end and 10 nt from 3’ end) using FASTX trimmer, whereas single end read files were filtered with the FASTX quality filter so that only reads where ≥ 70% of bases with a minimum quality score of 25 were retained (http://hannonlab.cshl.edu/fastx_toolkit/index.html). Reads were mapped to their corresponding genomes using TopHat2 version 2.0.9 (Kim et al., 2013). TopHat2 settings varied based on organism and SRA, and are listed in File S1. Transcripts were assembled using the Cufflinks2 app version 2.1.1 under settings listed in File S1 (Trapnell et al., 2010). TopHat2 and Cufflinks2 were executed on reads from each SRA file independently.

### Validation of lincRNA expression in vivo

RNA was extracted from 2-week seedlings and 4-week flower buds from Arabidopsis Col-0 using Trizol (ThermoFisher Life Sciences catalog # 15596018). These tissues and age at extraction most closely matched the experiments from which the RNA-seq data was obtained (Liu et al., 2012). cDNA was synthesized using SuperScript III (ThermoFisher Life Sciences catalog # 18080051) and 2μg of RNA as input. Primers were first validated by performing PCR with genomic DNA as template using GoTaq Green polymerase master mix (Promega catalog #M712) with 95°C for 3’ to denature, followed by 35 cycles of 95°C for 15”, 55°C for 30” and 72°C for 30” and a final extension step of 5’ at 72°C. Primers used are listed in File S2.

## Results

### An overview of lincRNA identification with Evolinc-I

#### Evolinc-I validation

After establishing a pipeline using the most commonly accepted parameters for defining a lincRNA (detailed in Materials and Methods), we wanted to know how well it removed unknown or novel protein-coding genes. For this, we tested Evolinc-I on a random set of 5,000 protein-coding transcripts selected from the TAIR10 annotation to determine the false discovery rate (FDR) (i.e., protein-coding transcripts erroneously classified as lincRNAs). ORFs for this test dataset of known protein-coding genes ranged from 303 to 4182 nts, with an average ORF length of 1131 nts (File S3). Because Evolinc is designed to automatically remove transcripts that map back to known genes, we removed these 5,000 genes from the reference genome annotation file, then generated a transcript assembly file from RNA-seq data where these 5,000 genes were known to be expressed. We fed the transcript assembly file to Evolinc-I. Out of 5,000 protein-coding genes, only 11 were identified as lincRNAs by Evolinc-I (0.22% FDR; indicated within File S3). Further investigation of these 11 erroneously identified lincRNAs revealed that they were predominantly low coverage transcripts with ORFs capable of producing polypeptides greater than 90, but less than 100 amino acids (aa). Moreover, low read coverage for these transcripts led to incomplete transcript assembly. These factors were responsible for the miss-annotation of these loci as lincRNAs. Importantly, these results indicate that read depth and transcript assembly settings impact lincRNA identification, a finding also noted by Cabilli et al. (2011). Therefore, exploring transcript assembly parameters may be necessary prior to running Evolinc-I. In sum, Evolinc-I has a low FDR that can be further reduced by increasing read per base coverage thresholds during transcript assembly as performed in Cabilli et al. (2011).

We determined the overlap of Evolinc predicted lincRNAs with previously published datasets from humans and Arabidopsis, following as closely as possible the methods published for each dataset. We first used Evolinc-I to identify lincRNAs from an RNA-seq dataset generated by Liu et al. (2012) in Arabidopsis (File S1). From nearly one billion reads generated from four different tissues (siliques, flowers, leaves, and roots), Liu et al. (2012) identified 278 lincRNAs (based on the TAIR9 reference genome annotation). Using the Liu et al. (2012) SRA data, we mapped RNA-seq reads and assembled transcripts with Tophat2 and Cufflinks2 in the DE. From these transcripts, Evolinc-I, identified 571 lincRNAs. We then reconciled the lincRNAs identified in Liu et al. (Liu-lincRNAs) and using Evolinc-I (Evolinc-lincRNAs), by identifying overlapping genomic coordinates for lincRNAs from the two datasets using the Bedtools suite (Quinlan and Hall, 2010). Of the 278 Liu-lincRNAs, 261 were also recovered by Evolinc-I (Table S1). Cufflinks failed to assemble the 17 unrecovered Liu-lincRNAs, due to low coverage, and thus differences in recovery for these loci reflect differences in the Cufflinks parameters employed.

The Arabidopsis genome reference has been updated since Liu et al. (2012), from TAIR9 to TAIR10 (Lamesch et al., 2012). We also ran Evolinc-I with the TAIR10 annotation and found that only 198 of the 261 Liu-lincRNAs were still considered intergenic (Figure 1B). The remaining 63 were reclassified as overlapping a known gene (either sense overlapping transcript, SOT, or antisense overlapping transcript, AOT). This highlights an important aspect of Evolinc-I. While Evolinc-I is able to identify long non-coding RNAs without a genome annotation, genome annotation quality can impact whether an lncRNA is considered intergenic versus AOT or SOT. In sum, 198 of the 571 lincRNAs identified by Evolinc-I correspond to a previously identified Liu-lincRNA (Figure 1B).

Of the 571 lincRNAs identified by Evolinc-I, 373 were not classified as lincRNAs by Liu et al. (2012). Evolinc-I removes transcripts that overlap with the 5’ and 3’ UTRs of a known gene, whereas Liu et al. (2012) removed transcripts that were within 500 bp of a known gene (Liu et al., 2012). This difference in the operational definition of intergenic space accounts for the omission of 197 Evolinc-lincRNAs from the Liu et al. (2012) lincRNA catalog. In addition, Evolinc-I removes transcripts with high similarity to transposable elements, but not tandem di- or trinucleotide repeats. We could see no biological reason for excluding these simple repeat containing transcripts, and in fact, transcripts with simple tandem repeats have been attributed to disease phenotypes and therefore might be of particular interest (Usdin, 2008). The inclusion of these transcripts accounts for 106 of the unique Evolinc-lincRNAs.

Finally, 70 of the 571 Evolinc-lincRNAs were entirely novel, and did not correspond to any known Liu-lincRNA or gene within the TAIR10 genome annotation. To determine whether these represented false positives or reflected the benefits of a stream-lined pipeline, we tested expression of a subset (*n* = 20) of single and multi-exon putative lincRNAs by RT-PCR using RNA extracted from two different tissues (seedlings and flowers, Figure S1A). We considered expression to be positive if we recovered a band in two different tissues or in the same tissue but from different biological replicates. We recovered evidence of expression for 18 of these putative lincRNAs out of 20 tested. Based on this we conclude that a majority of the 70 novel lincRNAs identified by Evolinc-I for Arabidopsis are likely to reflect bona-fide transcripts, and thus valid lincRNA candidates.

We next compared Evolinc-I against a well-annotated set of human lincRNAs characterized by Cabili et al. (2011). Cabili et al. (2011) used RNA-seq data from 24 different tissues and cell types, along with multiple selection criteria to identify a “gold standard” reference set of 4,662 lincRNAs. We assembled transcripts from RNA-seq data for seven of these tissues (File S1) using Cufflinks under the assembly parameters and read-per-base coverage cut-offs of Cabili et al. (2011) (see Materials and Methods). We then fed these transcripts to Evolinc-I. To directly compare Evolinc-I identified lincRNAs with the Cabili et al. (2011) reference dataset (Cabili-lincRNAs), we used the BED files generated by Evolinc-I to identify a subset of 368 multi-exon putative lincRNAs that were observed in at least two tissues (consistent with criteria employed in Cabili et al. [2011] when using a single transcript assembler). We then asked whether these 368 Evolinc-I lincRNAs were found in either the Cabili-lincRNAs, or the hg19 human reference annotation (UCSC). A total of 317 (88%) of the Evolinc-I lincRNAs matched known lincRNAs from the two annotation sources (Figure 1C).

We explored the 43 Evolinc-I lincRNAs that lacked a Cabili-lincRNA counterpart. The primary difference between the approach taken by Cabili et al. (2011) and Evolinc-I is the use of PhyloCSF tracks (Lin et al., 2011). PhyloCSF utilizes multiple sequence alignments of homologous loci to infer ORF conservation (represented as a positive score value), a strong indicator of protein-coding potential. The PhyloCSF scores for these 43 loci were either negative or absent, indicating that an ORF was not conserved (transcript identity and genomic coordinates shown in File S2) and that they were likely not discarded by the Cabili method due to ORF conservation. Instead, these 43 transcripts (12% of the 360 tested) appear to be bonafide lincRNAs, but will require further testing. In total, 88% of Evolinc-I identified lincRNAs agreed with reference datasets, whereas an additional 12% may reflect novel lincRNAs. Due to anticipated lack of sequence homology or simple lack of genome data that users may deal with, we did not include ORF conservation as a filtering step within Evolinc-I, but instead suggest users to perform a PhyloCSF or RNAcode (Washietl et al., 2011) step after homology exploration by Evolinc-II.

### Evolution of lincRNA loci with Evolinc-II

#### Evolinc-II validation

Evolinc-II is an automated and improved version of a workflow we previously used to determine the depth to which Liu-lincRNAs (Liu et al., 2012) were conserved in other species of the Brassicaceae (Nelson et al., 2016). The Evolinc-II workflow is outlined in Figure 2A. While most Liu-lincRNAs were restricted to Arabidopsis, or shared only by Arabidopsis and *A. lyrata*, 3% were conserved across the family, indicating that the lincRNA-encoding locus was present in the common ancestor of all Brassicaceae ~54 MYA (Beilstein et al., 2010). We used Evolinc-II to recapitulate our previous analysis in three ways. First, to provide statistical significance to our data, we randomly divided the 5,361 Liu-lincRNAs into 200-sequence groups prior to Evolinc-II analysis (*n* = 27; Figure 2B and Figure S2B). A separate comparison was performed by dividing the Liu-lincRNAs based upon chromosomal location of their encoding gene (*n* = 5). Following Evolinc-II’s comparative genomic analysis, we recovered similar percentages of sequence homologs for the Liu-lincRNA dataset regardless of to which of the 27 sequences groups they were assigned, or on which Arabidopsis chromosome they were located (Figure 2B and Figure S2C). Lastly, we used Evolinc-II to search for syntenic sequence homologs using the complete Liu-lincRNA dataset but querying with varying E-value cutoffs (E-20, E-15, E-10, E-05, and E-01). This analysis allowed us to test the ability of our reciprocal BLASTn approach to remove false positives driven by a low stringency approach (Figure 2B and Figure S2D). The increase in the number of sequence homologs retrieved by adjusting the E-value became insignificant at the last E-values tested (comparing recovery at E-05 vs. E-01; Figure S2D). Thus, Evolinc-II is a robust method for examining genomic, and, when the data are available, transcriptomic conservation of lincRNA loci.

In addition to identifying subsets of lincRNAs that are conserved across a user-defined set of species, Evolinc-II also highlights conserved regions within each query lincRNA. To demonstrate these features, we scanned through the Liu-lincRNA Evolinc-II summary statistics file (at 1E-10; File S4) to identify a conserved lincRNA. At1NC023160 is conserved as a single copy locus in eight of the ten species we examined. It was identified by Liu et al. (2012) based on both RNA-seq and tiling array data, as well as validated by Evolinc-I. During the comparative analyses, Evolinc-II generates a query-centric coordinate file that allows the user to visualize within a genome browser what regions of the query lincRNA are most conserved. Using this query-centric coordinate file, we examined the 332 nt At1NC023160 locus in the CoGe genome browser and determined that it was the 3’ end of the locus that was conserved (Figure 2C). To highlight the usefulness of this information, we then used the MAFFT multiple sequence alignment generated by Evolinc-II for At1NC023160 to perform structure prediction with RNAalifold (Figure S3A; (Lorenz et al., 2011)). As is often the case, the structural prediction based on the multiple sequence alignment was better supported than when examining the Arabidopsis lincRNA alone (Figure S3B and S3C). We feel that Evolinc-II’s ability to highlight conserved regions within a lincRNA locus will be useful for probing lincRNA function.

#### Using Evolinc-II to infer the evolution of the human telomerase RNA locus TERC

In addition to exploring the evolutionary history of a lincRNA catalog, Evolinc-II is an effective tool to infer the evolution of individual lincRNA loci. To showcase the insights Evolinc-II can provide for single lincRNA genes, we focused on the well-characterized human lincRNA, TERC. TERC is the RNA subunit of the ribonucleoprotein complex telomerase that is essential for chromosome end maintenance in stem cells, germ-line cells, and single-cell eukaryotes (Theimer and Feigon, 2006; Zhang et al., 2011; Blackburn and Collins, 2011). TERC is functionally conserved across almost all eukarya, but is highly sequence divergent. Building on work performed by Chen et al. (2000) we used Evolinc-II to examine the evolutionary history of the human TERC locus in 26 mammalian species that last shared a common ancestor between 100-130 MYA (Figure 3) (Glazko, 2003; Arnason et al., 2008).

Evolinc-II identified a human TERC sequence homolog in 23 of the 26 species examined (Figure 3; raw output shown in Figure S4). We were unable to identify a human TERC homolog in *Ornithoryhnchus anatinus* (platypus), representing the earliest diverging lineage within class Mammalia, using our search criteria. In addition, *Mus musculus* (mouse) and *Rattus norvegicus* (rat) were also lacking a human TERC homolog. However, close relatives of mouse and rat, such as *Ictidomys tridecemlineatus* (squirrel) and *Oryctolagus cuniculus* (rabbit) retained clear human TERC sequence homologs, suggesting that loss of the human TERC-like locus is restricted to the Muridae (mouse/rat family). This is in agreement with the previous identification of the mouse TERC, which exhibits much lower sequence similarity with the human TERC than do other mammals (Chen et al, 2000). All identified human TERC homologs also share synteny, suggesting similar evolutionary origins for this locus throughout mammals (Figure 3). Evolinc-II also identified lineage-specific duplication events for the human TERC-like locus in the orangutan, lemur, galago, cat and ferret genomes (Figure 3), similar to previous observations in pig and cow (Chen et al., 2000). In sum, Evolinc-II can be applied to both large and small datasets to uncover patterns of duplication, loss, and conservation across large phylogenetic distances.

**Figure 3.**
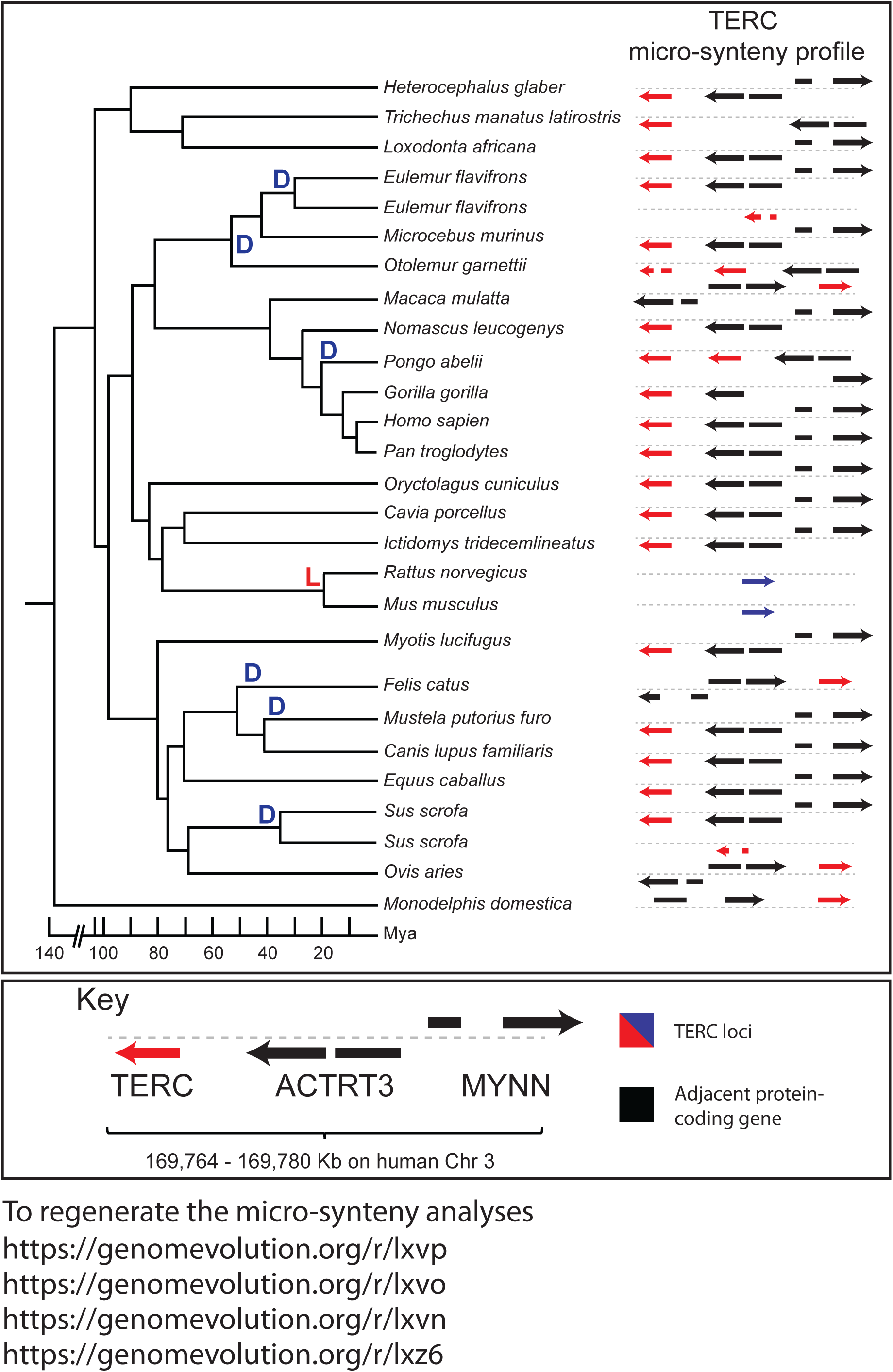
Evolinc-II analysis of the human TERC locus in mammals. Species tree of 26 species within class Mammalia with duplication (D) or loss (L) events hung on the tree (left). A micro-synteny profile is shown to the right for each species, showing the TERC locus in red, and adjacent protein-coding genes in black. Direction of each gene is indicated with arrows. The mouse and rat TERC loci are indicated by blue arrows to represent the poor sequence similarity between these two loci and human TERC. Divergence times are approximate and extracted from Arnason et al. (2008). A key is shown below, with gene names indicated. All pertinent links are shown below to regenerate micro-synteny analyses with CoGe (genomeevolution.org) for all species on
the tree.

## Discussion

### Rapid identification of lincRNAs using Evolinc-I

With Evolinc-I we set out to develop an automated pipeline for rapid lincRNA discovery from RNA-seq data. In addition to identification, Evolinc-I generates output files that put downstream analyses and data visualization into the hands of biologists, making it simpler for researchers to generate and explore lincRNAs. Evolinc-I makes use of standard lincRNA discovery criteria, and packages each step into easy-to-use applications within the CyVerse DE. In this way, Evolinc-I takes advantage of the cyberinfrastructure support of CyVerse (Merchant et al., 2016). One of the key advantages of combining Evolinc-I with cyberinfrastructure such as the CyVerse’s DE is the ability to combine various applications together in one streamlined workflow, and making the workflow easier to implement by interested researchers. For instance, a user can download an RNA-seq SRA file into their DE account, quickly process and map reads, assemble transcripts, and execute Evolinc-I. All of this occurs within the DE without downloading a single file or installing a program on a desktop computer.

We demonstrated the ability of Evolinc-I to identify lincRNAs from previously curated catalogs for plants and mammals. Note that we were able to account for all differences between results from Evolinc-I and the published studies, indicating that our pipeline is operating under definitions and filters currently used by the community. Moreover, because we have formalized the process by which annotations of genome data can be incorporated into the search strategy, Evolinc-I gives researchers the ability to easily explore the contributions of TEs, repetitive elements, or other user defined features to the prediction of lincRNA loci. Finally, we stress that this tool permits experiments to be repeated by researchers to compare the contribution of recently released annotations, or to repeat experiments from other groups. This latter point cannot be overemphasized as interest in lincRNAs grows.

### Examining evolutionary history and patterns of conservation of lincRNA loci using Evolinc-II

Evolinc-II is designed to perform a series of comparative genomic and transcriptomic analyses across an evolutionary timescale of the user’s choosing and on any number (1-1000s) of query lincRNAs. The computationally intensive nature of these analyses are ameliorated by once again taking advantage of CyVerse’s cyberinfrastructure. The analyses performed by Evolinc-II highlight conserved lincRNA loci, conserved regions within those loci, and overlap with transcripts in other species. We envision Evolinc-II being useful for both scientists attempting to identify functional regions of a lincRNA as well as those wanting to understand the pressures impacting lincRNA evolution.

In addition to highlighting large-scale lincRNA patterns of conservation, we also demonstrated how Evolinc-II can be used to examine the detailed evolutionary history of a single lincRNA, using the human TERC as a test-case. We performed an Evolinc-II analysis with human TERC on 26 genomes in the class Mammalia, 25, 14 of which were not examined by Chen et al. 2000. As expected, we recovered a human TERC-like locus in most mammals, as well as five previously unrecorded lineage-specific duplication events. Whether these duplicate TERC loci are expressed and interact with telomerase is unknown; if so they may represent potential regulatory molecules, similar to TER2 in Arabidopsis (Xu et al., 2015; Nelson and Shippen, 2015). We also determined that the human TERC-like locus was lost (or experienced an accelerated mutation rate relative to other mammals) in the common ancestor of mouse and rat, suggesting a novel TERC locus has been acquired in these species. The conservation of the TERC loci across mammals, characterized by rare evolutionary transitions such as that in mouse and rat, stands in stark contrast to the evolution of the telomerase RNA in Brassicaceae (Beilstein et al., 2012). Interestingly, mammalian TERCs appear to evolve more slowly than their plant counterparts, similar to the protein components of telomerase (Wyatt et al., 2010). These discoveries highlight the novel insights that can be uncovered using Evolinc-II on even well studied lincRNAs.

In summary, Evolinc streamlines lincRNA identification and evolutionary analysis. Given the wealth of RNA-seq data being uploaded on a daily basis to NCBI’s SRA, and the computing capabilities of CyVerse, we believe that Evolinc will prove to be tremendously useful. Combining these resources, Evolinc can uncover broad and fine-scale patterns in the way that lincRNAs evolve and ultimately help in linking lincRNAs to their function.

## Acknowledgements

We thank Evan Forsythe (University of Arizona) and Dr. Molly Megraw (Oregon State University) for thoughtful comments pertaining to Evolinc parameters. We thank the PaBeBaMo discussion group at the University of Arizona, and in particular Drs. David Baltrus, Rebecca Mosher, and Ravi Palanivelu, for critical comments on this work. We would also like to thank the CoGe group at the University of Arizona. We are very grateful for the cloud computing resources at CyVerse, and in particular the CyVerse DE group for their support in establishing Evolinc as a set of apps in the discovery environment. This work was supported by National Science Foundation Plant Genome Research Program Grant IOS 1444490 to M.A.B. and E.L. and NSF-MCB #1409251 to MAB.

---

**File S1** List of publically available genome and sequence files used, as well as conditions and results from TopHat and Cufflinks for each assembly.

**File S2** Evolinc-I output for all species from which lincRNAs were identified, as well as bed files for genome browser viewing, and primers used in RT-PCR verification of transcription of novel Arabidopsis lincRNAs. Also contains CoGe genome browser links to the novel lincRNAs identified.

**File S3** False-positive testing of Evolinc-I with Arabidopsis protein-coding genes.

**Figure S1.**
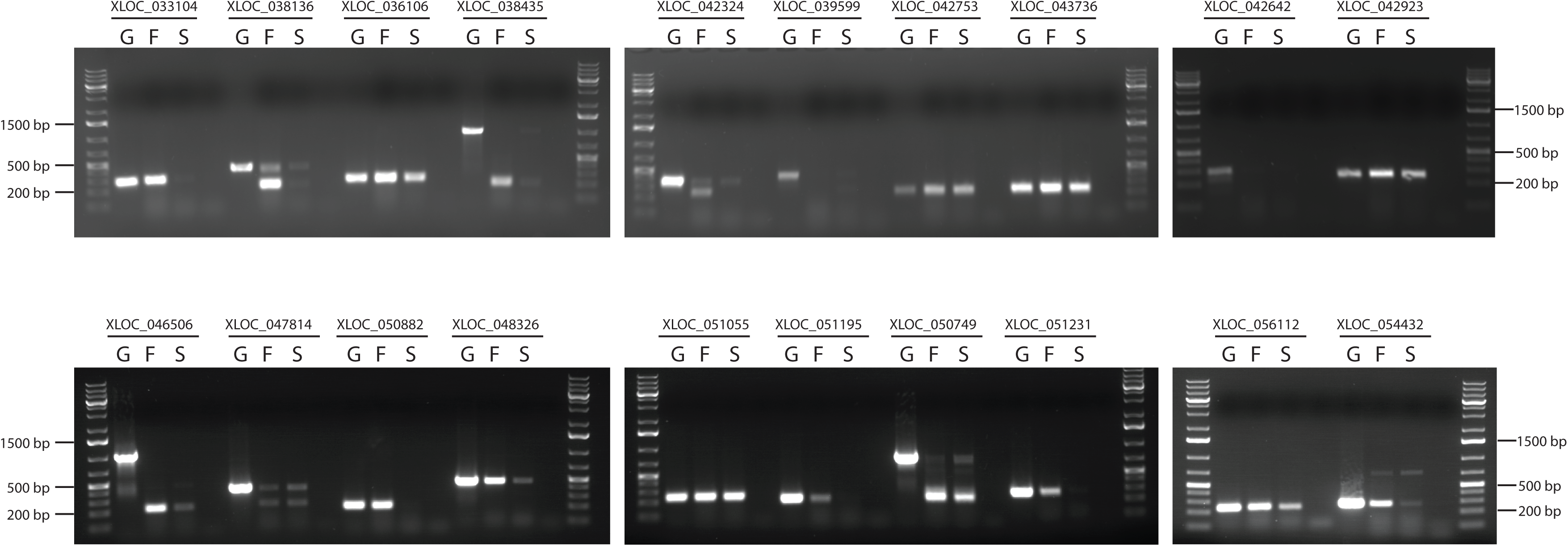
RT-PCR validation of lincRNAs identified in Arabidopsis by Evolinc-I. LincRNA IDs match those found in File S2. G = genomic DNA positive control. F = flower cDNA, S = seedling cDNA.

**Figure S2.**
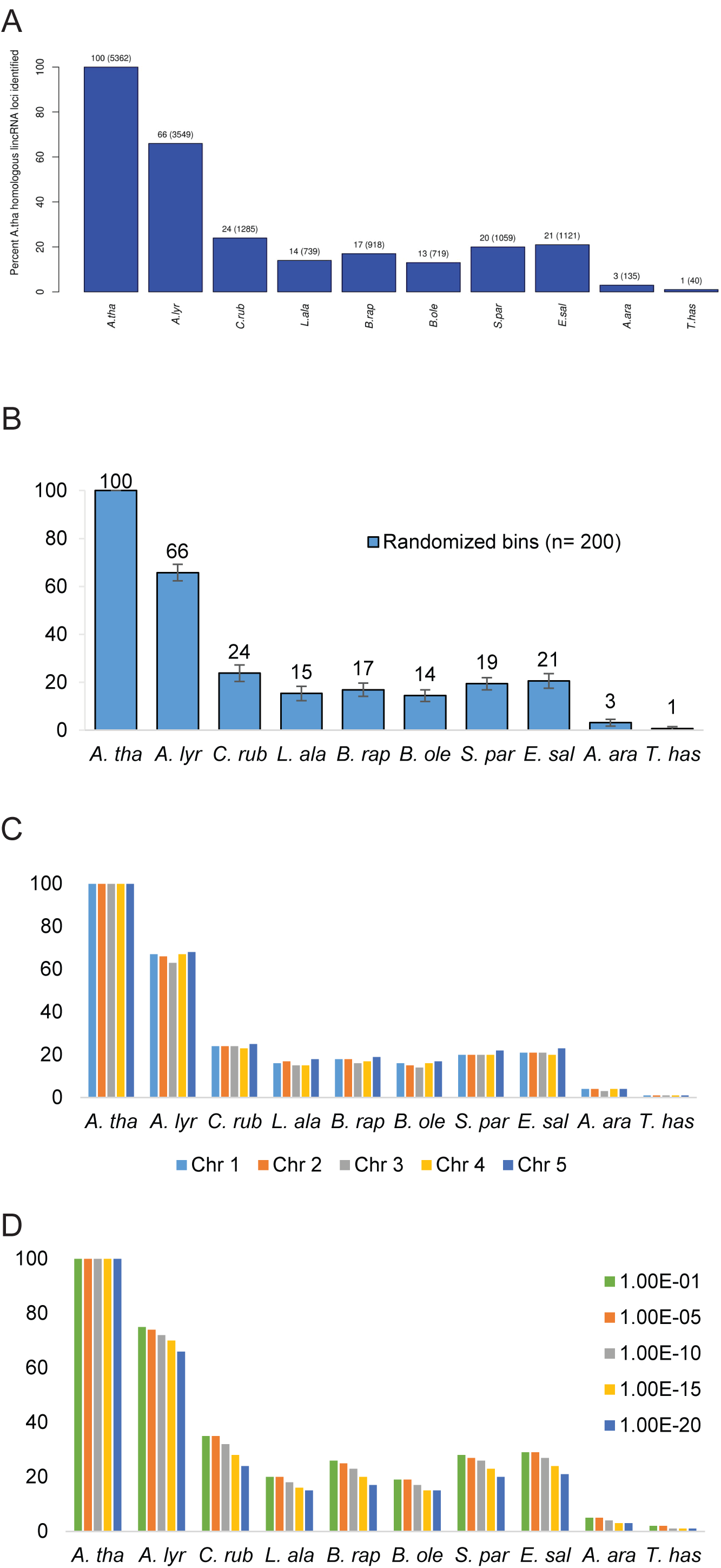
Examining conservation of Liu-lincRNAs in multiple ways with Evolinc-II. **(A)** Example of the type of bar graph produced by Evolinc-II, in this case for the Liu-lincRNAs at 1E-20. **(B)** Bar graph of level of lincRNA conservation observed when dividing the Liu-lincRNAs into 27 random bins of 200 lincRNAs each. Standard deviation is based on the difference seen between the 27 bins. **(C)** Bar graph depicting the level of lincRNA conservation seen when dividing the Liu-lincRNAs by Arabidopsis chromosome (E-cutoff value of 1E-20). **(D)** Bar graph demonstrating the level of conservation of the Liu-lincRNAs throughout Brassicaceae at different E-cutoff values.

**Figure S3.**
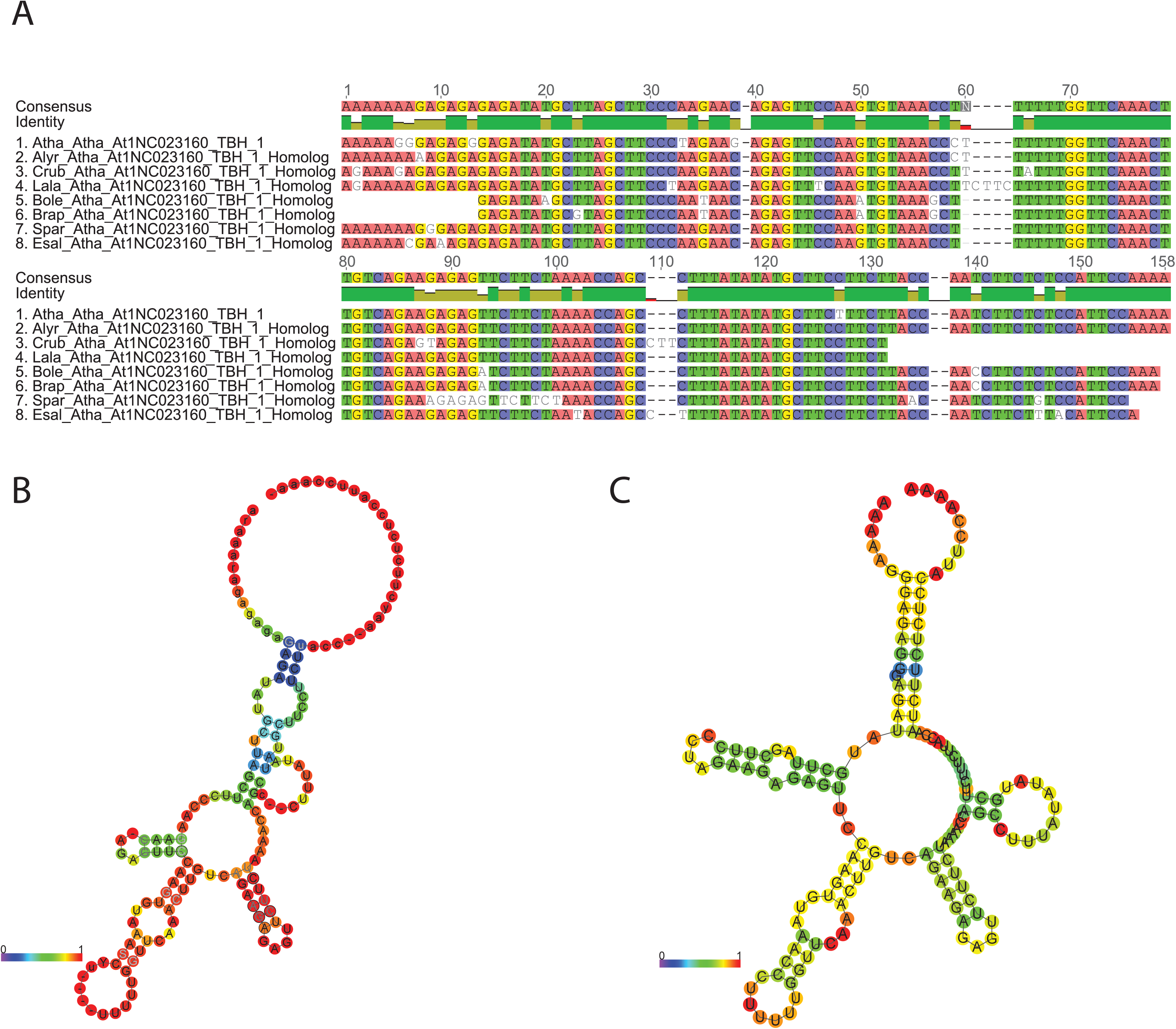
Using At1NC023160 to highlight the structural information that can be gleaned from Evolinc-II. **(A)** Multiple sequence alignment, generated by MAFFT and visualized within Geneious v7.1 (Kearse et al., 2012). Similar sequences are highlighted, with the consensus sequence shown on top. Nucleotide identity is shown below the consensus sequence, with green representing 100% identity across all sequences. **(B)** RNAalifold (Lorenz et al., 2011) consensus secondary structure prediction based on multiple sequence alignment in **(A)**. Base-pair probabilities are shown, with red being more probable and blue least probable. **(C)** RNAfold structure prediction based on the same region as in **(B)**, but limited to just the Arabidopsis sequence. Base-pair probabilities are shown as in **(B)**.

**Figure S4.**
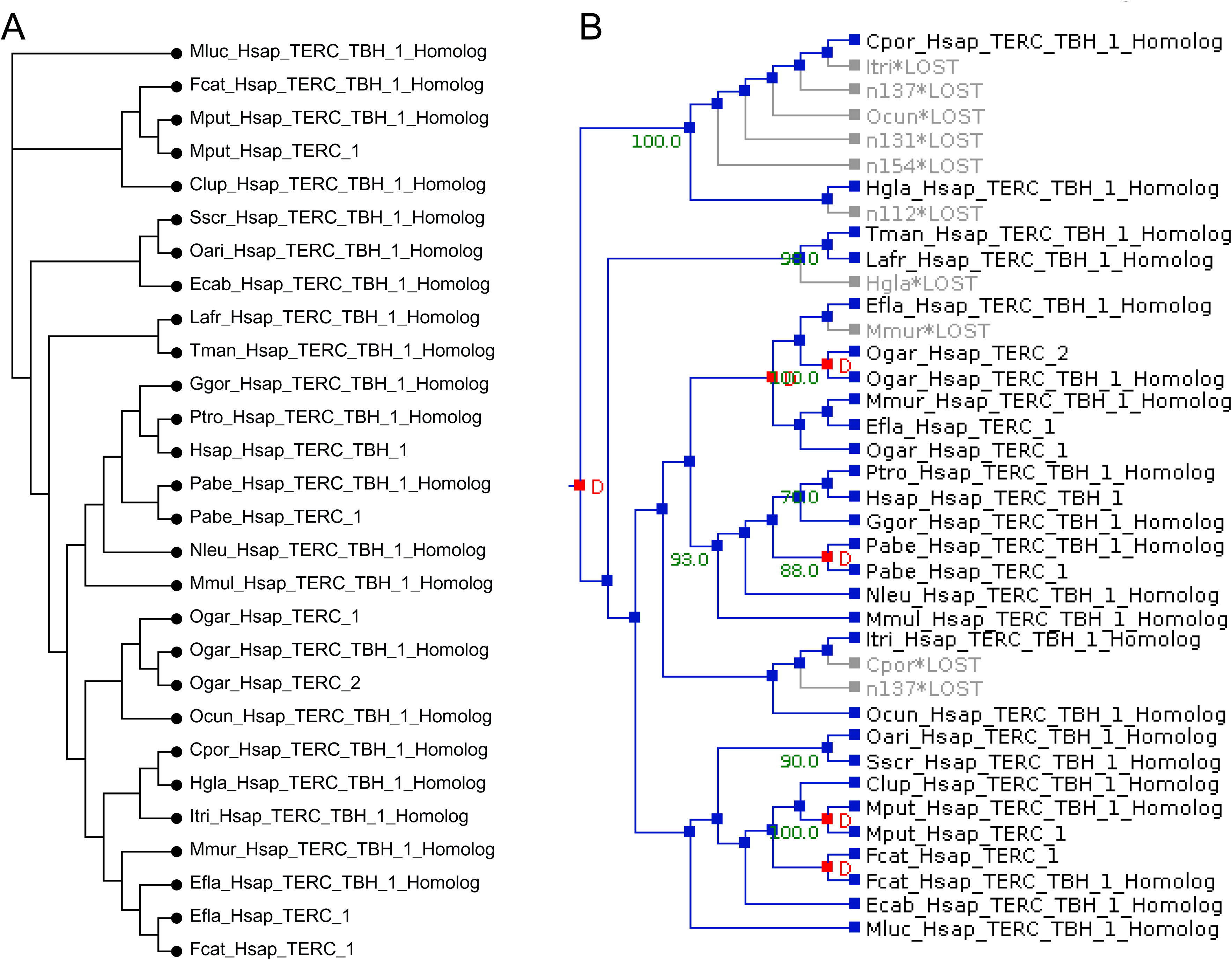
Raw phylogenetic output from Evolinc-II for TERC. **(A)** A gene tree for the TERC sequence homologs identified in each of the species shown. Sequences without “TBH” indicate paralogs. **(B)** Notung (Durand et al., 2006) reconciliation of the gene tree shown in (A) to the known species tree. Duplication (red “D”) and loss events (grey “LOST”) are shown. Support for duplication or loss events are indicated by the green numbers at the nodes that represent the predicted origin of those events.

**Table S1.** Percent similarity between transcripts identified following transcript assembly and lincRNA identification.

|  | <u>Identified transcription units overlapping between studies</u> |  | Percent similarity |
| --- | --- | --- | --- |
|  | Liu et al., 2012 | Present study |  |
| ATU | 30650 | 32911 | 107.3 |
| GATU | 370 | 352 | 95.1 |
| RCTU | 678 | 643 | 94.8 |
| lincRNA | 278 | 261 | 93.8 |

ATU = Annotated Transcription Unit
GATU = Gene Associated Transcription Unit
RCTU = Repeat Containing Transcription Unit
Nomenclature taken from Liu et al., 2012

